# Reduced alpha amplitudes predict perceptual suppression

**DOI:** 10.1101/2020.11.15.383562

**Authors:** Eva Poland, Aishwarya Bhonsle, Iris Steinmann, Melanie Wilke

## Abstract

The amplitude of prestimulus alpha oscillations over parieto-occipital cortex has been shown to predict visual detection of masked and threshold-level stimuli. Whether alpha activity similarly predicts target visibility in perceptual suppression paradigms, another type of illusion commonly used to investigate visual awareness, is presently unclear. Here, we examined prestimulus alpha activity in the electroencephalogram (EEG) of healthy participants in the context of a generalized flash suppression (GFS) task during which salient target stimuli are rendered subjectively invisible in a subset of trials following the onset of a full-field motion stimulus. Unlike for masking or threshold paradigms, alpha (8-12 Hz) amplitude prior to motion onset was significantly higher when targets remained subjectively visible compared to trials during which the targets became perceptually suppressed. Furthermore, individual prestimulus alpha amplitudes strongly correlated with the individual trial-to-trial variability quenching following motion stimulus onset, indicating that variability quenching in visual cortex is closely linked to prestimulus alpha activity. We conclude that predictive correlates of conscious perception derived from perceptual suppression paradigms differ substantially from those of obtained with “near threshold paradigms”, possibly reflecting the effectiveness of the suppressor stimulus.

## INTRODUCTION

Conscious perception is a constructive process which relies not only on physical input but also on the internal state of the brain. This is evidenced by visual illusions where conscious stimulus perception is intermittently interrupted, e.g. when stimuli are presented near-threshold and by so called ‘perceptual suppression’ paradigms where salient stimuli are erased from visual awareness ^1^. Perceptual suppression paradigms include motion-induced blindness, binocular rivalry and continuous or generalized flash suppression ^1–3^. In perceptual suppression (as opposed to near-threshold) paradigms, targets are well visible before they are rendered invisible by a competing stimulus. For example, in the generalized flash suppression (GFS) paradigm that we employ in the current study, a high-contrast target stimulus is shown close to central vision for several hundred milliseconds followed by the onset of a moving surround. The onset of the surround renders the target invisible in a subset of trials ^4^, with higher probabilities of target suppression with feature-specific preadaptation, spatial closeness as well as interocular discrepancy between target and surround. While the all-or-none fashion of target disappearance in GFS suggests the contribution of higher-order visual competition and selection processes, the target adaptation requirements underline the importance of pre-surround processes involving topographically organized (i.e. earlier) visual cortices. Roughly consistent with this interplay between low- and higher level contributions, neurophysiological experiments that employed GFS or related suppression paradigms have identified neural activity differences between visible and invisible trials in a wide range of brain areas and at different spatial and temporal scales. Specifically, electrophysiological studies in humans and monkeys found that reported periods of perceptual target suppression coincide with a decrease of low frequency power (~8-30 Hz) in occipito-parietal cortex, accompanied by spiking/gamma power changes in higher-order visual and prefrontal cortices ^5–11^. In contrast to the neural correlates of perceptual suppression states, the neural factors that *lead* to perceptual suppression in some trials but not in others remain unclear. Adopting the recent terminology of the consciousness problem, we learned about the neural correlates and/or consequences while lacking knowledge about the enabling, pre-requisite factors of flash suppression ^2,12,13^. This gap is likely due to the usage of paradigms where stimuli disappear spontaneously (e.g. MIB) and the common practice to normalize the data to the pre-stimulus (or pre-surround onset) baseline ^5,10,14^.

In the current study we employed GFS to ask whether upcoming perceptual suppression can be predicted from neural activity before the onset of the surround stimulus. We primarily focus on oscillatory power in the alpha frequency band, thought to reflect neural excitability states as evidenced in electrophysiological ^15–17^ and transcranial brain stimulation studies ^13,18^. Higher pre-stimulus alpha power has been linked to visibility in near-threshold paradigms, with higher alpha power serving as a predictor for impaired target detection and awareness / confidence reports^19–24^. Building on those studies, one could predict pre-surround alpha power to be higher in trials where targets become suppressed upon surround onset. On the other hand, if GFS suppression relies on biasing the target-surround competition toward the surround and/or represents a higher-level visual selection process, one might expect alpha power to be lower in suppression trials. The current study aimed to test these predictions.

The second aim of the study was to assess the contribution of trial-to-trial variability and its surround-induced decrease to perceptual suppression. Trial-to-trial variability and its decrease upon stimulus onset (‘quenching’) across a multitude of brain regions has been closely linked to visual perception and attention ^25–28^. Specifically, lower trial-to-trial variability predicts visual detection of stimuli presented at the perceptual threshold ^29^, and lower individual levels of variability quenching upon stimulus onset predict lower perceptual discrimination thresholds across subjects ^28,30^. Despite these functional similarities, trial-to-trial variability and alpha amplitude have been discussed in separate bodies of literature and only very recently, variability quenching has been related to the decrease in alpha/beta amplitude upon stimulus onset ^31^.

Thus, we employed GFS to evaluate whether alpha amplitudes prior to the onset of the surround stimulus predict perceptual suppression and investigated the relationship between prestimulus alpha amplitudes, perceptual suppression and variability quenching upon surround onset.

## RESULTS

We recorded 600 trials from 27 subjects performing a generalized flash suppression (GFS) task ^4^, a visual illusion by which salient target stimuli can be rendered subjectively invisible after the onset of a random dot motion (RDM) stimulus (**Figure 1A**). Following central fixation, subjects were presented with two target stimuli consisting of two red disks, one in the left and right visual hemifield for 2 seconds. Then, the RDM stimulus was added to the presentation for 2 seconds. After the end of stimulus presentation, subjects were asked to report whether the right, the left, both or neither target had perceptually disappeared. Perceptual outcomes are summarized in **Figure 1B**.

**Figure 1.**
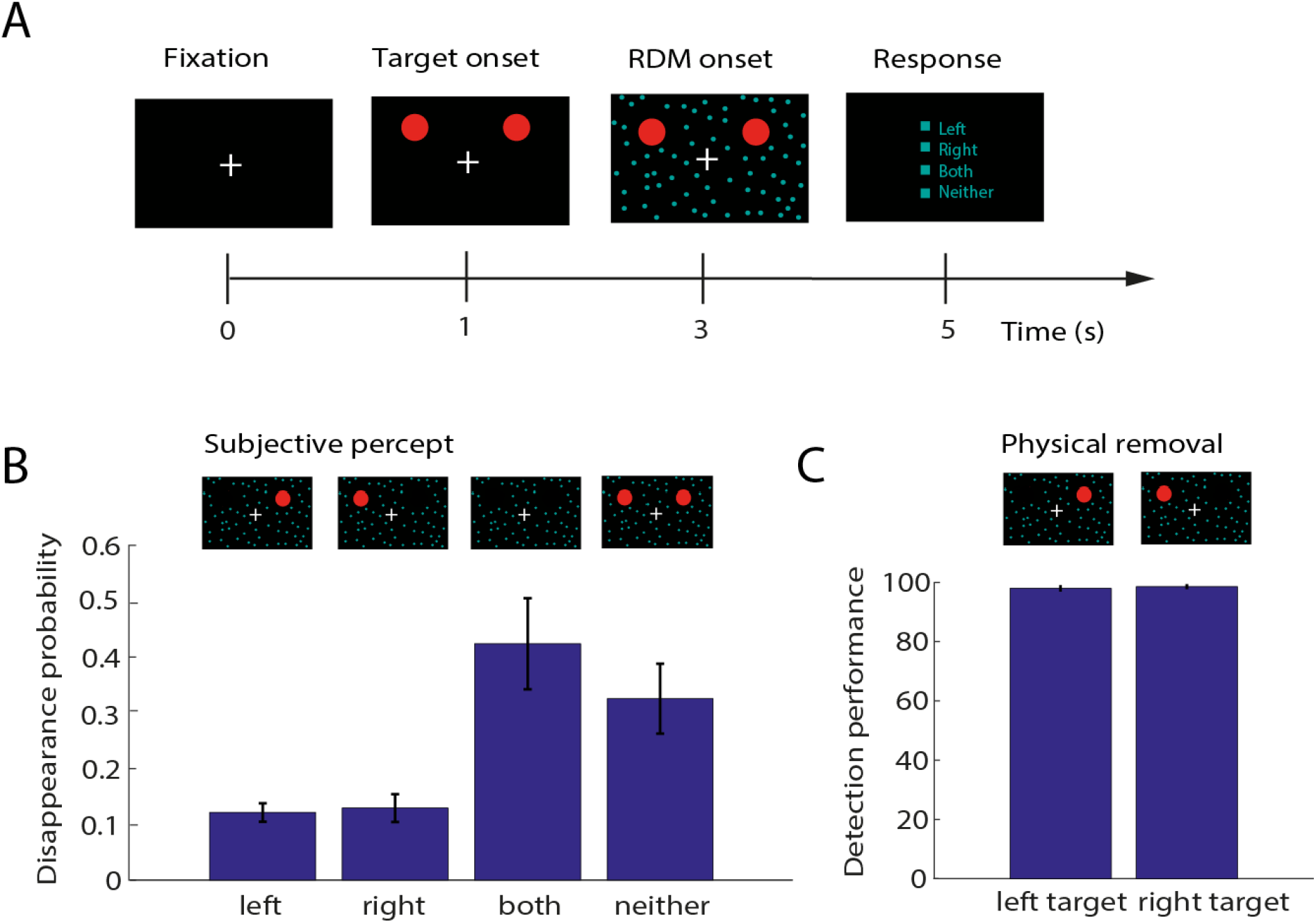
Generalized flash suppression (GFS) paradigm and perceptual results. **(A)** Each experimental trial began with central fixation, followed by the onset of two salient red target stimuli in the upper left and right visual hemifield. After the target had been presented for 2 seconds, a random dot motion (RDM) stimulus was shown, resulting in the disappearance of one or both target stimuli in a subset of trials. After each trial, subjects reported whether the left, the right, both or neither target had disappeared by selecting one of the options with a mouse click. Stereoscopy was achieved using red-green anaglyphic glasses. **(B)** Mean probabilities of the left, right, both or neither target disappearing for all 27 subjects. Corresponding subjective percepts are illustrated above the respective bars. **(C)** Percentage of correct reports of physical removal of the left or the right target during the control conditions. In all plots, error bars indicate +/− 1 SEM across subjects.

Generally, stimuli were more often suppressed or remained subjectively visible together rather than disappearing individually, although unilateral disappearances of only the left or the right target occurred in a small portion of trials (left target: 0.12 +/− SD 0.9, right target: 0.13 +/− SD 0.7, both targets: 0.42 +/− SD 0.26, neither target: 0.33 +/− SD 0.27). As a control, each session further included a total of 120 catch trials during which either the right or the left target stimulus was physically removed from screen. Accuracy in detecting these physical removals was very high in all subjects (98% +/− SD 3% for left target removals and 99% +/− SD 2% for right target removals, **Figure 1C**), indicating that subjects performed the task correctly and were attentive throughout the experimental session.

We then evaluated whether parieto-occipital prestimulus alpha activity reflected the subjective visibility of the targets. The time course of mean alpha amplitude across subjects around the time of the onset of the RDM stimulus for trials in which both targets had disappeared (‘invisible’) and trials in which neither target had disappeared (‘visible’) is shown in **Figure 2A**. Due to the long target adaption requirement of the GFS (2 seconds), we considered a time window prior to the onset of the RDM stimulus as the prestimulus interval rather than a time window preceding the target. Comparing the average alpha amplitude of all parieto-occipital electrodes in the second preceding the onset of the RDM stimulus, we observed a significant difference between trials during which the targets later became suppressed and trials during which they remained visible (Wilcoxon signed-rank test, p = 0.0039, N = 27, mean alpha amplitude visible: 4.41 +/− SD 2.36, mean alpha amplitude invisible: 4.23 +/− SD 2.56). We did not observe any differences between perceptual conditions in alpha amplitude post RDM stimulus (poststimulus window, Wilcoxon signed-rank test, p = 0.94). The visible - invisible difference in prestimulus alpha amplitude was positive in most subjects regardless of the mean alpha amplitude across perceptual conditions, suggesting that prestimulus alpha amplitudes were consistently lower on trials during which the targets were perceptually suppressed independent of the individual mean amplitude (**Figure 2B**). **Figure 2C** shows the FFT of the second prior to RDM stimulus onset across subjects for visible and invisible trials. The mean individual alpha frequency (IAF) across subjects was 9.89 Hz +/− SD 1.37, ranging from 7 to 13 Hz. Comparing power specifically at the subjects IAF, we found IAF power to be significantly higher on visible compared to invisible trials (Wilcoxon signed-rank test, p = 0.008, N = 27), indicating that the effect was in fact due to an amplitude difference.

**Figure 2.**
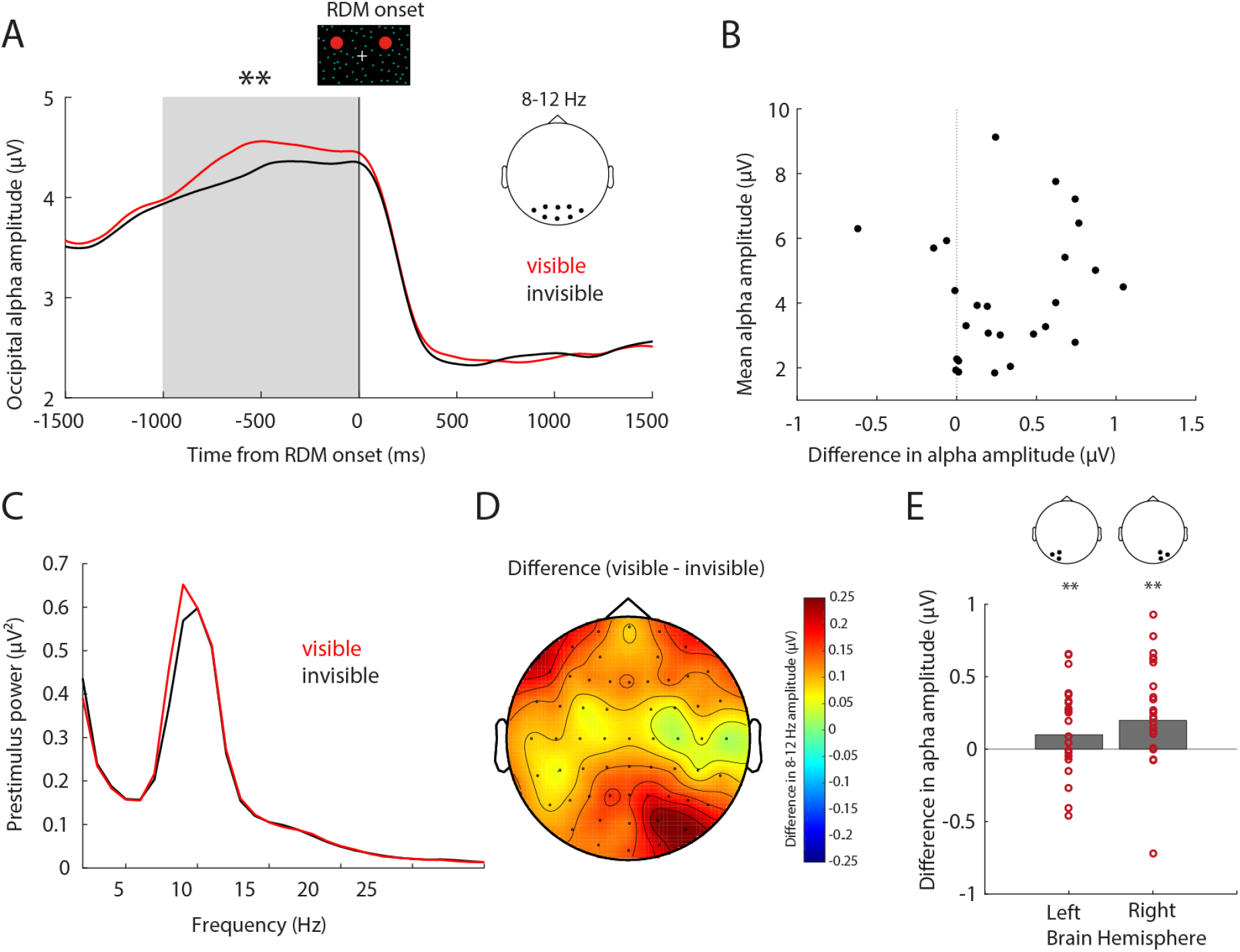
Prestimulus alpha amplitude as a function of perceptual suppression. **(A)** Average time course of alpha band (8-12Hz) amplitudes for trials in which both targets remained subjectively visible (red, ‘neither’ condition) and trials in which both targets were perceptually suppressed (black, ‘both’ condition, N = 27). The zero mark denotes the onset of the RDM stimulus following which the targets were perceptually suppressed. Data represent the mean of all parietal-occipital electrodes. ** denotes a significant group difference in alpha amplitude as assessed by Wilcoxon signed-rank test (p < 0.01). **(B)** Difference in parieto-occipital alpha (8-12 Hz) amplitude between visible and invisible conditions (visible – invisible) in the second preceding RDM onset for individual subjects as a function of the individual average 8-12 Hz amplitude in the second preceding RDM onset across perceptual conditions, outlier corrected > 2SD, N = 26. **(C)** Fast Fourier Transform (FFT) of the second preceding the RDM stimulus for visible and invisible trials across subjects. **(D)** Topography of the visible – invisible difference in 8-12 Hz amplitude in the second prior to RDM onset, N = 27. **(E)** Difference in parieto-occipital alpha (8-12 Hz) amplitude between visible and invisible conditions (visible – invisible) in the second preceding RDM onset, separated by left (electrodes O1, PO3, PO7) and right hemisphere (RH, electrodes O2, PO4, PO8) separately. Each dot represents the average of a single subject (outlier corrected > 2SD, N = 25). ** denotes a significant alpha amplitude difference between visible and invisible trials as assessed by Wilcoxon signed-rank test (p < 0.01), separated by hemisphere.

We further determined the individual peak alpha frequency during the second prior to RDM onset for visible and invisible trials separately and found them to not statistically differ (visible 9.85 Hz +/− SD 1.32, invisible 9.92 Hz +/− SD 1.36, Wilcoxon signed-rank test p = 0.69, N = 27). The topography of the visible – invisible difference in alpha amplitude across all 63 electrodes (**Figure 2D**) suggested that the effect was most prominent in right parieto-occipital cortex. The spatio-temporal clustering analysis on 100 ms windows we conducted to identify visibility-related prestimulus differences independent of the a priori selected electrodes and the 1 second time interval revealed a significant difference between visible and invisible conditions (cluster-level statistic = 42.35, p = 0.04, N = 27) that was most prominent between 600-400 ms prior to RDM onset and at right occipital and frontal electrodes (**Supplementary Information, Figure S1**). Comparing alpha amplitudes during the second preceding RDM onset between visible and invisible trials in each hemisphere separately (**Figure 2E**), we found that both hemispheres contributed to the effect (Wilcoxon signed-rank tests, left hemisphere p = 0.007, right hemisphere p = 0.002, N = 27). Upon examining unilateral disappearances as a function of hemisphere, we found no significant differences between trials in which the left target and trials in which the right target disappeared individually (**Supplementary Information, Tables S1, S2**).

Although previously discussed in separate bodies of literature, findings characterizing the role of prestimulus alpha activity in visual perception bear many similarities to those reported for neural variability across trials. In particular, the magnitude of the variability reduction upon stimulus onset has previously been shown to be a reproducible trait closely linked to individual perceptual abilities ^28^, and very recently, to reflect stimulus-induced decreases in alpha power ^31^. Inspecting time courses of trial-to-trial variance and alpha amplitudes of individual subjects, we noted that both measures closely co-varied (**Figure 3A**). As expected from previous studies ^25,28,30,32^, variability across trials significantly decreased with the onset of the RDM stimulus in parieto-occipital cortex (relative variance, Wilcoxon signed-rank test, p = 0.00009, N = 27). Likely due to the ongoing visual stimulation by the dynamic RDM pattern, trial-to-trial variability remained at a constant lower level during the 2 second stimulus presentation interval. Subjects notably differed in the degree of variability quenching defined as the mean relative variance in a stable interval 500 – 1500 ms post stimulus capturing the variance decrease (**Figure 3B**).

**Figure 3.**
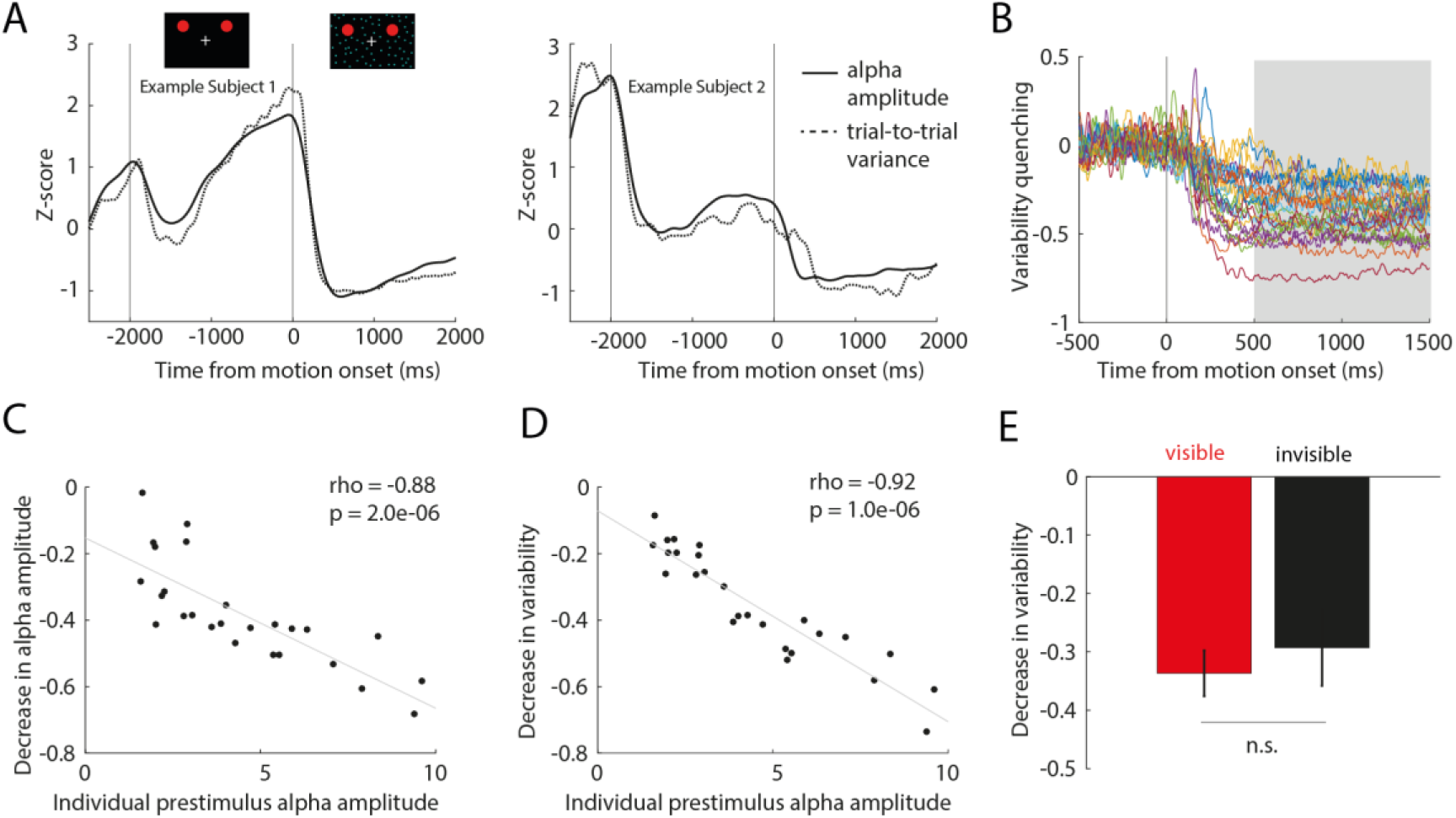
Quenching of trial-to-trial variability and relation to prestimulus alpha amplitude. **(A)** Average variance across trials (z-score, dotted line) and alpha amplitude (z-score, black line) over the course of experimental trials for two example subjects. **(B)** Individual relative variance per subject (outlier corrected >2SD, N = 26) and post stimulus time window used to define the individual degree of variability quenching. **(C)** Relation of decrease in alpha amplitude to individual alpha amplitude in the prestimulus interval, N = 26. Each dot represents one subject. Spearman’s rank correlation coefficient and corresponding p-value. The gray line represents the least-squares fit. **(D)** Relation of the individual degree of variability quenching to individual alpha amplitude in the prestimulus interval, N = 26. Spearman’s rank correlation coefficient and corresponding p-value. The gray line shows the least-squares fit. **(E)** Mean change in variance with RDM stimulus onset (trial-to-trail variability quenching) for visible and invisible trials across subjects (N = 26). Error bars indicate +/− 1 SEM.

Consistent with previous findings ^31^, we found the individual magnitude of variability quenching to be significantly correlated with the decrease in alpha amplitude following the onset of the RDM stimulus (Spearman’s rank correlation coefficient, rho = 0.87, p = 0.000002, N = 26). Based on the similarities of previous findings associating both trial-to-trial variability quenching as well as prestimulus alpha amplitude with subsequent perceptual performance, we wondered whether the individual decrease in trial-to-trial variance as well as alpha amplitude was intrinsically related to the individual magnitude of ongoing alpha activity prior to stimulus onset. We found both the decrease in alpha amplitude (Spearman’s rank correlation coefficient, rho = −0.88, p = 0.000002, N = 26) as well as the decrease in trial-to-trial variability (Spearman’s rank correlation coefficient, rho = −0.92, p = 0.000001, N = 26) to be significantly negatively correlated to the individual prestimulus alpha amplitude in the second preceding RDM stimulus onset (**Figure 3C-D**). Bandpass filtering the data to remove the influence of the different physiological frequency bands ^31^ revealed that oscillations in the alpha frequency constituted the largest contribution to prestimulus variance and also had the largest effect on variability quenching (**Supplementary Information S3, Figure S2**).

Importantly, neither the magnitude of variance in the second preceding RDM onset nor during the stable period of RDM stimulus presentation differed significantly between invisible and visible trials (Wilcoxon signed-rank tests, prestimulus p = 0.70, poststimulus p = 0.35). The comparison of variability quenching between visible and invisible trials revealed no significant difference between perceptual states (Wilcoxon signed-rank tests, p = 0.75, N = 26, **Figure 3E**), suggesting that variability quenching itself did not differentiate between perceptual outcomes.

## DISCUSSION

In the current study, we employed generalized flash suppression (GFS) to compare parieto-occipital prestimulus alpha activity of human EEG data between physically identical trials during which visual stimuli underwent perceptual suppression or remained visible. In contrast to previous studies that used visual masking or perceptual threshold paradigms and have found prestimulus alpha power to be reduced for stimuli that are consciously perceived ^19–21,23^, we observed significantly lower alpha amplitudes prior to perceptual suppression compared to trials during which the targets remained visible. One possible explanation for our result is that the varying perceptual outcomes of GFS, despite identical physical stimulation, are due to spontaneous fluctuations in attentional state. Attentional modulations of neural activity serve to preferentially process behaviourally relevant stimuli while inhibiting the processing of competing distractors, and previous research has found alpha activity to be reduced for attended compared to unattended stimuli ^33–35^. Decreased alpha activity has been shown to reflect a state of increased neural excitability ^36,37^ and has been associated with distractor suppression ^33,38,39^. In our case, reduced prestimulus alpha activity prior to RDM onset would thus predict enhanced processing of the surrounding motion stimulus, rendering it more effective in suppressing GFS targets. Unlike previous studies investigating the relationship between prestimulus alpha activity and visual awareness in near-threshold paradigms, we examined alpha activity prior to the onset of the motion stimulus rather than preceding the target itself. It is thus conceivable that in paradigms in which a target stimulus never reaches awareness, target visibility may depend on neural excitability prior to target onset, whereas in our case, increased neural excitability prior to the upcoming motion stimulus predicts its effectiveness to actively suppress an already consciously perceived target stimulus from awareness.

Given that in the current study we were not able to distinguish between alpha modulations pertaining to the target and the suppressor stimulus presented in the same visual hemifield, another possibility is that the reduced alpha activity we observed prior to perceptual suppression via GFS reflects increased attentiveness towards the already present target stimulus, which subsequently facilitated its subjective disappearance. Behavioral evidence for this interpretation comes from previous studies using motion-induced blindness (MIB), a related perceptual suppression paradigm where salient stimuli spontaneously disappear due to a moving surround pattern^40^. Specifically, using MIB in combination with an attentional cueing task that required subjects to report hue changes in one of two target stimuli, Schölvinck and Rees demonstrated that directing spatial attention to a MIB target counterintuitively increases the probability of its disappearance compared to the unattended target ^41^. Previous psychophysical GFS experiments showed that the effectiveness of the surround stimulus strongly depends on pre-surround target adaptation ^4,6^. GFS shares this pre-adaptation requirement with other perceptual suppression illusions such as MIB and binocular rivalry flash suppression (BRFS), while being considerably shorter than Troxler fading or filling-in illusions ^1,42^. It is thus conceivable that the amplitude of alpha oscillations before motion onset could have influenced the probability of perceptual suppression by affecting target adaptation, which has previously been suggested to increase the relative salience of upcoming novel stimuli ^43^.

In previous intracranial recordings in monkeys, Wilke and colleagues showed that perceptual suppression was reflected in an alpha power decrease of local field potentials (LFP) at target-responsive sites in striate and extrastriate visual cortex ^5,6^. This modulation started after RDM onset and corresponded to the estimated illusory disappearance time. Consistent modulations of intracranial spectral amplitude by perceptual suppression have also been observed in humans ^10^. In the current study, we did not observe any visibility-related differences in alpha amplitude post RDM stimulus, but it is possible that such modulations are too local to be evident in parieto-occipital EEG alpha activity that is largely driven by the full-field motion stimulus. In addition to methodological differences between intracranial electrophysiological recordings and those of EEG sum potentials that pool activity from neural populations representing both target and motion stimulus, there could be perceptual report-related factors that account for the absence of significant post RDM effects in our study ^3^. Specifically, in contrast to the previous GFS studies in monkeys ^5,14^, in the present study subjects reported their percept only after passively viewing the RDM stimulus for 2 seconds rather than indicating target disappearances immediately. It is thus conceivable that the delayed report resulted in reduced task engagement during stimulus presentation. It is also noteworthy that we asked subjects to report the occurrence of subjective disappearances only after RDM presentation regardless of whether the target had reappeared, and we thus cannot assume stable target suppression in the invisible trials.

In addition to lower alpha amplitudes prior to perceptual suppression, we observed a strong correlation between individual prestimulus alpha amplitude and the degree by which neural variability across trials declined with RDM onset. The magnitude of variability quenching has previously been shown to be an individual trait associated with individual perceptual abilities ^28,30^ and recently, to reflect the amplitude of neural oscillations in the alpha/beta band that similarly decreases with visual stimulation ^31^. Here, we provide evidence that these measures depend on the level of prestimulus alpha activity. Given that stimulus-induced decreases of trial-to-trial variability and alpha amplitude strongly correlate and both are related to the magnitude of alpha amplitudes prior to stimulus onset, it could be argued that the perceptual relevance of the relative rather than the absolute variance across trials ^28^ may be ultimately rooted in prestimulus alpha oscillations. In our case, only prestimulus alpha amplitude differentiated between perceptual outcomes while variability quenching did not, but more research is needed to understand the exact relationship between trial-to-trial variability and alpha amplitude and their consequences for perception.

While it seems evident that perceptual suppression critically depends on cortical state as indexed by prestimulus alpha oscillations in posterior cortex, several potential factors suppressing salient target stimuli from visual awareness remain unclear. Future studies specifically manipulating spatial attention could address the effect of attention on perceptual suppression and prestimulus alpha activity directly, as well as develop experimental designs that allow for the differentiation between modulations of alpha activity pertaining to the target and the motion stimulus. From the present study, we can conclude that reduced alpha activity predicts perceptual suppression, likely facilitating stronger processing of the upcoming motion stimulus and its effectiveness in suppressing the target.

## METHODS

### Subjects

A total of 35 healthy subjects participated in the current study. Of those, 4 subjects were excluded from the analysis due to general exclusion criteria (one due to red-green color blindness, one due to the subject reporting no subjective disappearances under generalized flash suppression (GFS), one due to <75% accuracy in the physical removal condition and one due to a technical error with the EEG recording setup). For the analysis of general visibility effects we compared trials in which both targets had disappeared (‘invisible’) and trials during which neither target disappeared (‘visible’). We required a minimum of 20 trials in each perceptual condition, resulting in a cohort of 27 subjects (12 male / 15 female, 14 left handers / 13 right handers, between 18 and 50 years of age). The mean number of visible trials was 159 +/− SD 133, the mean number of invisible trials was 215 +/− SD 146. All subjects gave informed consent and were paid for their participation in the study.

### Experimental procedure

Data were recorded in a single session lasting 3 to 4 hours in total, including EEG preparation, experiment and breaks. To exclude color blindness, subjects judged a set of 20 Ishihara plates of which 19 correct identifications were required for inclusion in the study ^44^. Prior to the experiment subjects performed 10 to 20 practice trials in order to familiarize themselves with the generalized flash suppression (GFS) task ^4^. We carefully instructed subjects to indicate target disappearances based on whether the target stimuli had disappeared completely, regardless whether perceptual suppression persisted until the end of the trial. Following EEG preparation, subjects completed 6 experimental blocks, each lasting approximately 10 minutes. Subjects were seated in front of a 60 × 34 cm computer screen and placed their head on a chin rest with an eye to screen distance of 70 cm. During the experiment, lights in the recording room were turned off and additional curtains were used to protect the subjects from extraneous light. Between blocks subjects had the opportunity to take brief breaks.

Stereoscopy was achieved using anaglyphic glasses. To control for individual differences in eye dominance, the positions of the red and green filters were interchanged with each configuration being used for 3 of the 6 experimental blocks in varied order to equate target presentations in the dominant and non-dominant eyes. All experimental procedures were approved by the ethics committee of the University Medicine Göttingen (UMG, Germany).

### Stimuli and task

Stimuli were programmed and presented in Matlab 2015b (The MathWorks Inc., Natick MA, USA) using Psychtoolbox-3 ^45,46^. Each trial began with central fixation for 1 second (**Figure 1A**). Subjects were then presented with two salient red target stimuli (size 3° of visual angle) in the left and right visual hemifield (7° of visual angle horizontal distance from center, 3° of visual angle vertical distance from center). The targets were only presented to one eye by means of anaglyphic red-green glasses. After 2 seconds, a random dot motion (RDM) pattern consisting of 2000 green dots moving at a speed of 10°/second was shown to the respective other eye for 2 seconds, which resulted in the subjective disappearance of one or both target stimuli in a subset of trials. Following the end of stimulus presentation, subjects were prompted to report their perception, that is the subjective disappearance of both, neither, the right or the left target stimulus, with a mouse click using their preferred hand equivalent to their handedness. In addition to the experimental trials (600 trials), each subject performed 120 control trials intermixed with the experimental trials in which either the left or the right target stimulus was physically removed following the onset of the RDM pattern.

### EEG acquisition and preprocessing

EEG activity was recorded from 63 electrodes without ECG signal (64-Channel Standard BrainCap, Brain Products, Gilching, Germany) using BrainVision Recorder (Brain Products, Gilching, Germany). Electrode impedances were kept below 20 kΩ throughout the experiment. EEG data were preprocessed and analysed using the Fieldtrip toolbox ^47^ and custom-written software in Matlab 2015b (The MathWorks Inc., Natick MA, USA). The data were recorded at a sampling rate of 1000 Hz and bandpass filtered between 0.1 – 250 Hz. Trials containing muscle artefacts, jumps or clipping artefacts were identified automatically and rejected following visual inspection. An independent component analysis (ICA) was performed to identify eye movement artefacts and eye movement related components were removed. The data were then re-referenced to a common reference.

### Prestimulus alpha amplitude

For the analysis of alpha amplitudes, the data of the parieto-occipital electrodes O1, O2, Oz, POz, PO3, PO4, PO7 and PO8 were bandpass filtered at 8-12 Hz with a 4^th^ order Butterworth filter and subsequently Hilbert transformed. We then considered the absolute values of the Hilbert transform, equivalent to the envelope of the filtered signal, in a time window spanning the second prior to the onset of the RDM stimulus (prestimulus window) pooled over the parieto-occipital electrodes for statistical analysis of prestimulus alpha amplitude. The selected electrodes cover occipital cortex and were chosen based on our expectation of possible modulations occurring in the visual system, as well as for consistency with previous research that investigated prestimulus alpha power within the same set of electrodes ^20^. For the main comparison between target visibility outcomes, we considered trials in which subjects had reported both targets to disappear (‘invisible’) and trials in which both targets had been reported to remain visible (‘visible’). In order to determine the subjects’ individual alpha frequency (IAF), we additionally calculated the Fast Fourier Transform (FFT) of the prestimulus time window for all parieto-occipital electrodes across perceptual outcomes between 1 and 30 Hz at a resolution of 1 Hz and identified the peak frequency between 5 and 15 Hz in each subject. We then calculated equivalent FFTs for visible and invisible trials separately and compared prestimulus FFT power at the IAF that was determined for each subject between perceptual outcomes. For topographical representation we calculated the average absolute value of the 8-12 Hz filtered Hilbert transform in the second prior to RDM onset for each channel separately. The difference between visible and invisible conditions was then calculated by subtracting the average prestimulus alpha amplitude of invisible trials from the mean prestimulus alpha amplitude of visible trials in each subject. For the comparison of alpha amplitudes in the left and right hemisphere we omitted the central electrodes Oz and POz of the initial selection and considered the lateral electrodes O1, PO3 and PO7 as left and O2, PO4 and PO8 as right hemispheric ROIs respectively. For the comparison of prestimulus alpha amplitude to stimulus-induced changes in trial-to-trial variability we computed the average prestimulus alpha amplitude per subject over all experimental trials regardless of their perceptual outcome. Due to the fact that most variables were not normally distributed, we applied non-parametric statistical methods throughout analyses and Wilcoxon signed-rank tests were used for within-subject comparisons.

### Trial-to-trial variability

In order to assess variability across trials, we first calculated the variance across trials for the signal of the same parieto-occipital electrodes O1, O2, Oz, POz, PO3, PO4, PO7 and PO8 for each 1 ms time point of each trial across all experimental trials for each subject. We then computed the relative variance as the percent change in variance with stimulus onset from a baseline 500 to 0 ms prior to the onset of the RDM stimulus. The degree of variability quenching for individual subjects was then determined as the average relative variance in a time window from 500 ms to 1500 ms post RDM stimulus, which best covered the variance decline (poststimulus window). We calculated the non-parametric Spearman’s rank correlation coefficient between the individual degree of variability quenching and the individual decrease in alpha amplitude based on the same baseline and post RDM stimulus window. Similarly, we calculated the Spearman’s rank correlation between the individual magnitude of prestimulus alpha amplitude across all experimental trials and the individual degree of variability quenching as well as the decrease in alpha amplitude with stimulus onset in order to determine whether the decreases were linked to prestimulus activity. To assess the perceptual relevance of stimulus-induced decreases of trial-to-trial variability we compared its change with motion onset using the same previously used baseline (−500 - 0 ms) and poststimulus window (500 - 1500 ms) between visible and invisible trials. For illustration of individual time courses of alpha amplitude and trial-to-trial variance, we z-scored both measures by subtracting the mean and dividing by the standard deviation according to z = (x – μ) / σ.

## ACKNOLEDGMENTS

This research was supported by the Hermann and Lily Schilling Foundation and the Leibniz Science Campus Primate Cognition. We thank Carsten Schmidt-Samoa and Severin Heumüller for help with the data acquisition software.

## AUTHOR CONTRIBUTIONS

E.P. and M.W. developed the study concept. E.P. and M.W. developed the task. A.B. and E.P. collected the data. E.P. analysed the data. I.S. provided analysis scripts. E.P., I.S. and M.W. interpreted the data and conceptually contributed to the analysis process. E.P. and M.W. drafted the manuscript. All authors provided critical comments during the manuscript writing process and approved the final version of the manuscript.

## DATA AVAILABILITY

The datasets acquired and analysed during the current study are available from the corresponding author on request.

