## Supplementary for "Reduced alpha amplitudes predict perceptual suppression"

### Supplementary Information S1: Cluster-permutation test

In addition to the analysis of the average alpha amplitude during the second preceding RDM onset, we performed a spatio-temporal clustering analysis allowing us to identify statistically significant differences in prestimulus alpha amplitude between visible and invisible conditions independent of the a priori selected electrodes and the 1 second time interval. To this end we divided the second prior to RDM onset into 100 ms windows. A cluster-level statistic was calculated based on these time windows of interest and all 63 sensors, requiring a minimum number of two neighbouring channels and consecutive time points. In order to obtain cluster-corrected p-values we employed a Monte Carlo permutation with 500 iterations. The cluster-permutation test revealed a significant difference between visible and invisible conditions (cluster-level statistic = 42.35, $p = 0.04$, $N = 27$) that was most prominent between 600-400 ms prior to RDM onset in and right occipital and frontal electrodes.

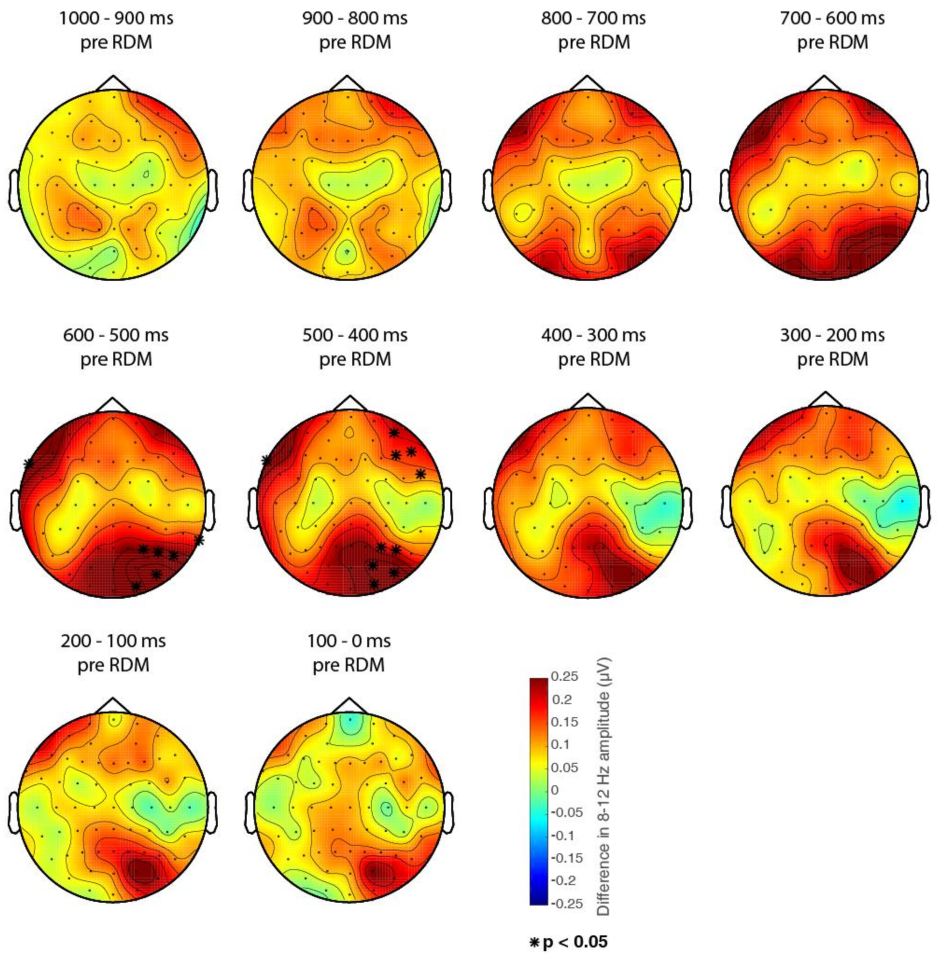


**Figure S1. Clustering analysis.** Scalp topographies of the visible –invisible difference in alpha amplitude (8 –12 Hz) for 100 ms windows spanning the second prior to RDM stimulus onset. Spatio-temporal clusters showing a significant difference between the two perceptual conditions are indicated by asterisks ($p < 0.05$).

### Supplementary Information S2: Unilateral target disappearances

**Supplementary Table S1** summarizes mean alpha amplitudes in the second prior to RDM stimulus onset for trials in which only either the left or the right target stimulus disappeared, separated by left and right hemisphere ROIs. Since previous studies investigating the lateralization of alpha activity have focused on more parietal sensors[1], we additionally compared unilateral target disappearances between left and right hemispheric ROIs that included more parietal electrodes (**Table S2**), but also did not observe a significant difference between left and right unilateral conditions when considering parietal sensors.

| | **Left hemisphere (O1 / PO3 / PO7)** | **Right hemisphere (O2 / PO4 / PO8)** |
|---|---|---|
| **Unilateral left disappearances** | 4.32 +/- SD 2.69 | 4.47 +/- SD 2.69 |
| **Unilateral right disappearances** | 4.15 +/- SD 2.29 | 4.40 +/- SD 2.43 |
| **p value** | 0.63 | 0.79 |

**Table S1.** Mean alpha (8-12 Hz) amplitudes in the second preceding RDM onset and standard deviations in the left and right hemisphere for unilateral target disappearances. Significance between unilateral left and unilateral right target disappearances as determined by Wilcoxon signed-rank tests, N = 21.

| | **Left hemisphere (PO3 / P5 / P7)** | **Right hemisphere (PO4 / P6 / P8)** |
|---|---|---|
| **Unilateral left disappearances** | 3.89 +/- SD 2.12 | 4.16 +/- SD 2.32 |
| **Unilateral right disappearances** | 3.76 +/- SD 1.93 | 4.12 +/- SD 2.20 |
| **p value** | 0.50 | 0.71 |

**Table S2.** Mean alpha (8-12 Hz) amplitudes in the second preceding RDM onset and standard deviations for unilateral target disappearances in more parietal left and right hemispheric electrodes. Significance between unilateral left and unilateral right target disappearances as determined by Wilcoxon signed-rank tests, N = 21.

### Supplementary Information S3: Frequency contributions to variability quenching

Finally, we sought to identify the contributions of different physiological frequency bands to the overall decrease in variability. To this end we first bandpass filtered the data between 4-90 Hz as a reference. For better comparability with the 4-90 Hz broadband signal (black, **Figure S2A**), we chose an indirect approach and removed the influence of the different physiological frequency bands one at a time by band-stop in the theta (4-7 Hz), alpha (8-12 Hz), beta (13-30 Hz), low gamma (31-50 Hz) and high gamma (51-90 Hz) bands using a 4$^{th}$ order Butterworth filter[2]. We then computed the variance across trials of all perceptual conditions for the filtered signals (**Figure S2A**) as well as the relative variance as percent change from baseline to post RDM stimulus interval. Oscillations in the alpha frequency (red) constituted the largest contribution to prestimulus variance and also had the largest effect on variability quenching. Significant changes in magnitude of quenching following band-stop filtering occurred for the removal of theta (44% of the full 4-90 Hz signal to 47%, p = 4.1000e-05), alpha (44% to 24%, p = 9.3386e-06) and low gamma (44% to 50%, p = 0.0059) frequencies, but not for beta (44% to 44%, p =0.39) and higher gamma (44% to 49%, p = 0.02, Bonferroni corrected p = 0.01). **Figure S2B** shows the average prestimulus amplitudes for the considered frequency bands in the second preceding RDM onset, scaling to the respective contributions to prestimulus variance.

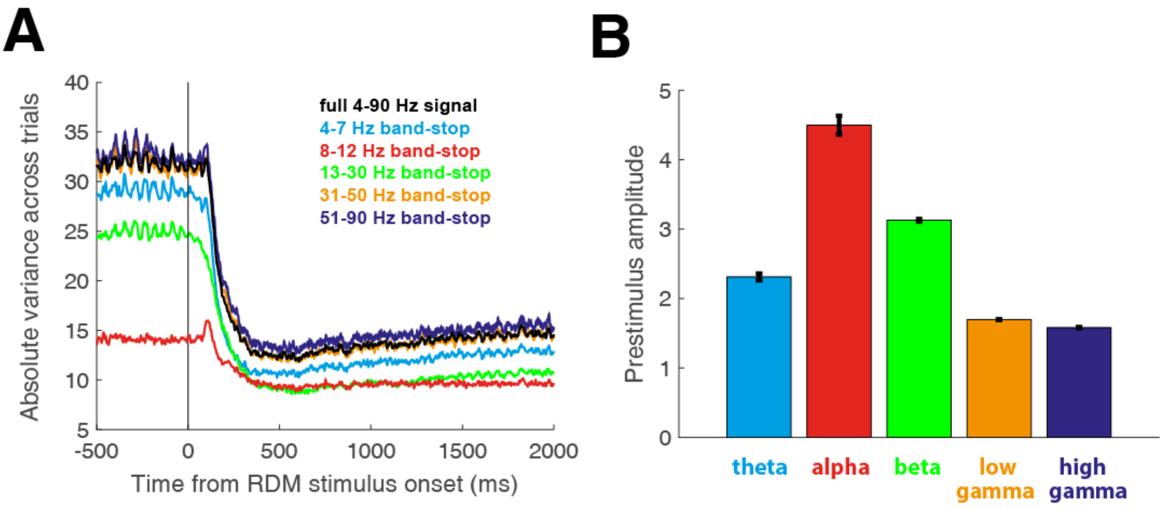


**Figure S2. Contributions of different frequency bands to variability quenching in visual cortex. A)** Average absolute variance around RDM stimulus onset for band-stop filtered broad-band signals from which the influence of different physiological frequency bands has been removed (N=27). **B)** Average prestimulus amplitude of oscillations in the corresponding physiological frequency bands theta (4-7 Hz, blue), alpha (8-12 Hz, red), beta (13-30Hz, green) as well as low (31-50 Hz, orange) and high gamma (51-90 Hz, purple). Error bars indicate +/- 1 SEM.
